# Diversification of pre-mating behaviors through temporal reordering of components

**DOI:** 10.1101/2025.10.22.683896

**Authors:** Elijah Carroll, Taisuke Kanao, Nathan Lo, Nobuaki Mizumoto

**Affiliations:** Department of Entomology and Plant Pathology, Auburn University, 301 Funchess Hall, Auburn, Alabama 36849 USA; Faculty of Science, Yamagata University, Yamagata, Japan; School of Life and Environmental Sciences, The University of Sydney, Camperdown, New South Wales 2006, Australia; Okinawa Institute of Science & Technology Graduate University, Onna-son, Okinawa, 904-0495 Japan

**Keywords:** Order dependence, modularity, precopulatory behavior, functional integration, social insects

## Abstract

Many biological processes, including mating behavior, are structured as ordered sequences in which each step enables the next. Diversity of such processes usually arises through the addition or loss of components rather than through changes in their order, since reordering can disrupt functional integration. Here we show that reordering of the steps associated with termite nuptial pairing can generate behavioral diversity. In most termites, dispersed adults shed their wings before initiating tandem pairing, as wings interfere with communication between females and males. However, we found that in the Australian termite, *Microcerotermes nervosus*, tandem running occurs prior to wing shedding, and the act of tandem running itself facilitates dealation. Video tracking revealed that wing removal improves movement coordination, and comparative analysis of sympatric species showed that *M. nervosus* exhibits much higher tandem stability than others. These results suggest that enhanced movement coordination of this species may have potentiated step reordering. Our findings demonstrate that even functionally conserved sequences can be reorganized and that overcoming such constraints can open novel pathways for behavioral diversification.

## Introduction

From embryonic development to nest construction, biological processes are typically structured as ordered sequences in which each step sets the condition for the next [1–3]. Because downstream events depend on the proper completion of upstream ones, the process cannot reach a functional endpoint when the order is disrupted. This is especially true for goal-oriented behaviors, where early actions create the physiological or informational context needed for later actions to succeed [2,4,5]. For example, in many *Drosophila* species, males perform a series of courtship behaviors in a fixed order, and experimental disruption of this order severely reduces female receptivity [6,7]. In such systems, although the details of each step may vary across species, their temporal order is usually conserved.

Termite precopulatory behavior exemplifies this rule particularly within Neoisoptera, where the behavioral sequence is functionally linked and conserved across species. The sequence begins when winged adults (alates) seasonally emerge from their colonies and disperse through flight to increase outbreeding opportunities [8,9]. After landing, alates shed their wings, and these unwinged termites (dealates) engage in a random search for a mating partner by walking [10,11]. Upon encountering each other, a female and male form a tandem running pair, with the male following closely behind the female. This allows the pair to maintain contact and undertake nest searching as a coordinated reproductive unit [10]. The pair then finds a nest site, excavates the site together, and initiates reproduction. During this process, each step creates the conditions for the next, making re-ordering of the steps unlikely. However, our understanding of this sequence is biased by observations from a set of economically important termite species [10,12,13], leaving open the question of cryptic diversity being present across other taxa.

Wing shedding (dealation) is a candidate step for testing whether sequences involved in nest searching and reproduction are modifiable. Following dispersal, alates voluntarily shed their wings by applying upward pressure to a basal suture on their wings using their abdomen [14]. This is a functional and essential behavior for termites. On the ground, wings hinder their mobility and can increase the probability of being captured by predators [14]. Moreover, wings interfere with finding a partner and coordination during tandem running [13,15,16]. Yet, previous documentations suggest that there is considerable variation in the timing of wing-shedding among species; the presence of conspecifics can facilitate wing-shedding in some species, while isolation or flight alone is sufficient to trigger dealation in others [14,17–20]. This diversity suggests that dealation acts as a modular component that may shift in position within the sequence rather than being fixed to a single position, generating novel behavioral patterns despite strong functional integration.

Here we report a previously undescribed precopulatory sequence in the termite species *Microcerotermes nervosus*. Unlike other termites, we found that this species initiates tandem running prior to shedding wings, and tandem running behavior facilitates wing shedding (Fig. 1D). Our video tracking demonstrates that dealation increases tandem running stability, and that *M. nervosus* exhibits greater tandem stability than other sympatric species. Our results demonstrate that enhanced movement coordination enabled the re-ordering of a functionally integrated behavioral sequence, contributing to the generation of new forms of behavioral diversity, otherwise conserved across species.

**Figure 1.**
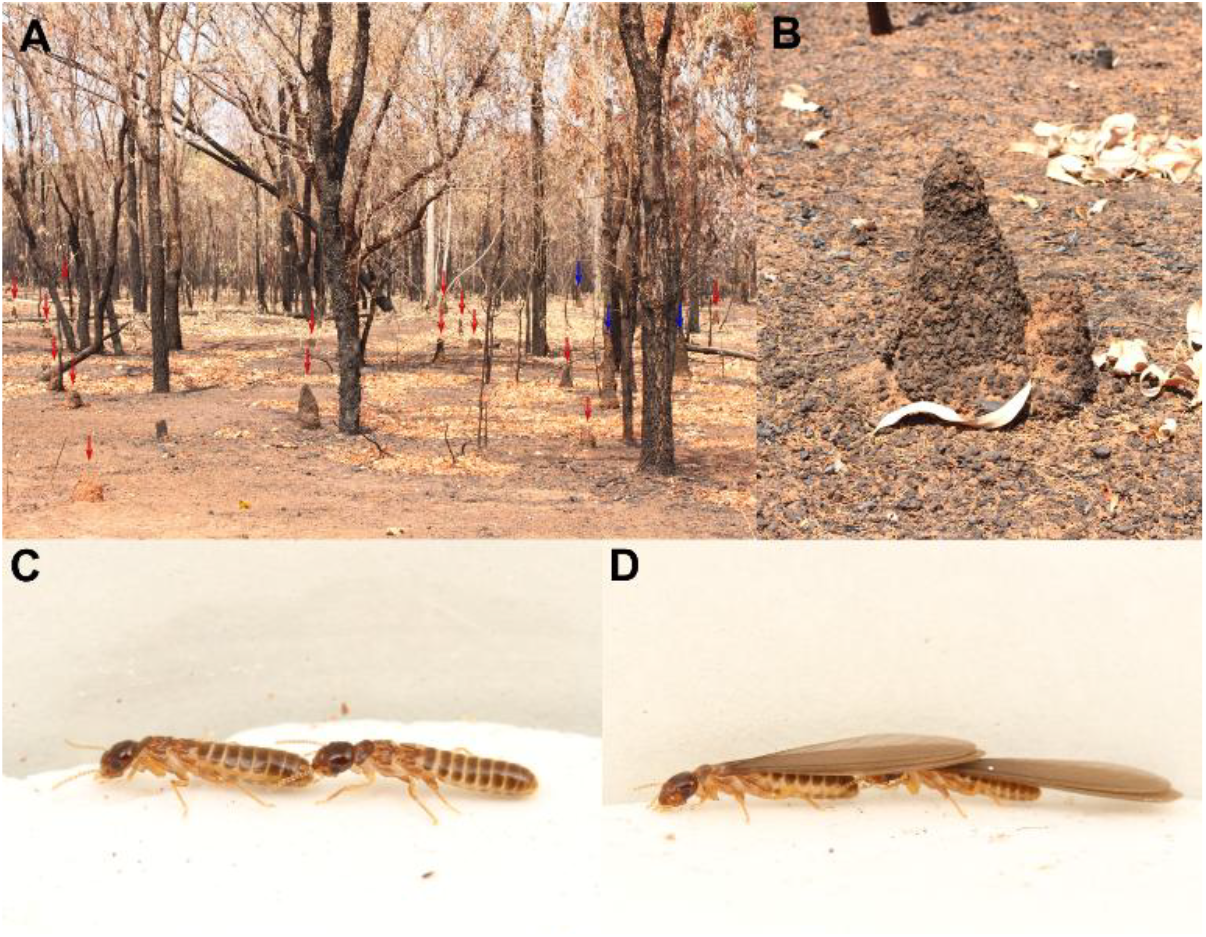
The primary study species, *Microcerotermes nervosus*. (A) Natural habitat of *M. nervosus* in the northern Australian tropical savanna. Red arrows indicate mounds of *M. nervosus*, while blue arrows indicate those of other species. (B) A closer look at the mound. Alates are aggregated at the top of the mound at the end of the dry season, waiting for the dispersal flight. (C-D) Unwinged- and winged-tandem running. Females are always leaders, while males are followers. Note that the wings of *M. nervosus* are not morphologically unique (e.g., length) compared to other species.

## Materials and Methods

### M. nervosus field collection and Behavioral Observations

We collected five colonies (A-E) of *M. nervosus* in Darwin and Howard Springs, Northern Territory, Australia, on Oct 28th, 29th, and Nov 1st, 2023. These dates were just before the rainy season (we detected the first rain on Nov 6th in 2023). Alates of *M. nervosus* swarm after the rain, and thus, many alates in the nest aggregated at the top of the mound at the end of the dry season to prepare for swarming (Fig. 1A & B). We cut the tip of the mound to collect alates with nests. After bringing back the nests, we sprayed them with water and disturbed them by opening them with axes and knives. Alates came out of the nests and flew to disperse. We collected these alates, separated by sex, and kept them with wet paper towels. All observations were performed within 12 hours of the flight. We isolated alates (or dealates) with wet filter paper for 30 minutes before the following behavioral observations.

To investigate the effect of the wings on tandem pairing in *M. nervosus*, we observed behavioral interactions between a winged female and a de-winged male. We focused on female wing status because females lead tandem running, and their wings can physically interfere with movement coordination. We introduced a pair to an observation arena, consisting of a petri dish (φ = 90 mm) filled with moistened plaster with a lid. We recorded the pair for 15 minutes at 30 frames per second with a video camera (HC-X1500-K, Panasonic) located vertically above the arena. All these videos were cropped to only include the dish arena, with both height and width being 2,000 px. Males started following their partner females with wings within a few seconds after introduction into the arena. During tandem runs, some females shed their wings upon separation and resumed the tandem run after re-encounter, while others slowed down and shed their wings

while actively engaging in tandem running. We obtained the videos of tandem runs led by the same females before and after wing shedding (Fig. 1C & D). We also observed the behavior of single isolated females with wings to identify the effect of the male partner on wing shedding behaviors. In total, we obtained the videos of 35 pairs (A: 12 replicates, B: 3 replicates, C: 9 replicates, D: 10 replicates, and E: 1 replicate), and 25 isolated females (A: 8 replicates, C: 8 replicates, D: 8 replicates, and E: 1 replicate). Before each observation, the plaster surface of the arena was wetted and polished to provide a clean surface.

We extracted the coordinates of moving termites from each video using either the video-tracking system, UMATracker [21] and Social LEAP Estimates Animal Poses (SLEAP) v 1.3.3 [22]. For a single alate female or a pair consisting of a female alate and a male dealate, we used UMATracker to obtain the trajectory of the body center. On the other hand, we used SLEAP for tracking females and males after dealation. This is because wings removed from the alate remained in the arena, which prevents a background-subtraction-based approach for movement tracking in UMATracker. Alternatively, SLEAP is based on the deep-learning framework, which is robust against noise created by wings. Note that, in order to reduce the time required for manual labeling, we did not use SLEAP for winged individuals, because such individuals have a different appearance from dealates. If females did not shed their wings in the presence of a conspecific male, termite trajectories were not tracked using SLEAP.

Six body parts were labeled to form a skeleton for pose estimation using SLEAP: head, tips of antennae (left and right), pronotum, body center (metathorax-abdomen junction), and the posterior tip of the abdomen. Among them, we used the midpoint between the head and the posterior tip of the abdomen for comparison with the tandem run with wings. Termite body lengths were calculated as the sum of two linear distances: from the head to the body center and from the body center to the posterior tip of the abdomen, averaged over all frames in each video. A model was initialized and trained from scratch to obtain trajectories. We labeled 48 frames in total for termite pairs. To infer termite tracks across frames, top-down and centroid networks were trained within an encoder-decoder (U-net) architecture with an EfficientNet-B0 encoder pretrained on ImageNet. The decoder consisted of four up-sampling blocks, each starting with 128 filters. Augmentation was performed only by rotating frames (from −180 to 180 degrees). Tracking was performed using the “simple” option with the Intersection over Union (IOU) similarity method and the Hungarian matching method. Misdetection of body parts was covered by interpolating the outputs with linear methods, followed by smoothing with a median filter (k =5). Pose estimation data from SLEAP were converted to HDF5 files for further analysis.

### Trajectory analysis

All videos were downsampled to 5 frames per second and scaled from pixels to mm (2,000 px = arena size) prior to analysis. To compare tandem running behaviors between pairs with alate and de-alate pairs, we automatically determined whether pairs were in tandem or not based on the distance between the body centers of termite pairs. We assumed that two individuals were tandem running when the distance was less than 1.6 body lengths; when the distance was greater than or equal to 1.6 body lengths, they were considered to be in separation [23,24]. We did not count short interactions (< 2 sec) as tandem running, nor short separations (< 2 sec) as separations. After smoothing, separations happened only after the first tandem running, i.e., pairs that exceeded this distance prior to the first tandem event were not counted as separated. To test whether wing shedding influenced the stability of coordinated movement, we compared tandem duration and separation duration for the same pairs before and after female dealation, where we removed the

data of females that did not shed their wings during the recording period. We analyzed tandem and separation duration using mixed-effects Cox proportional hazards models with the coxme() function in the ‘coxme’ package in R [25,26]. In each model, female wing status (winged vs. wingless) was included as a fixed effect, and video name nested within colony ID as a random effect structure to account for repeated measures within pairs. Model significance was assessed using Wald χ^2^ tests from the Anova() function in the “car” package [27]. In addition, we tested the effect of the male conspecific presence on the duration of female wing-shedding, using a similar mixed-effects Cox model as above.

We also determined movement speed by calculating the displacements between two successive frames for each female and male and then averaging these displacements. We compared movement speeds between tandem pairs before and after female wing shedding, and between isolated females and alate pairs, using a linear mixed model (LMM) using the lmer() function in the ‘lme4’ package in R [28], with colony included as a random effect. Model significance was assessed using Wald χ^2^ tests. If a female in the presence of a male did not shed her wings during the recording period, the data were removed from the analysis.

To test whether wing shedding affects the consistency of female movement speed during tandem, we modeled frame-to-frame acceleration with a linear mixed-effects framework that allows group-specific residual variances by wing status. For each focal female, we computed instantaneous acceleration every 0.2 seconds during tandem running. We fit two mixed models with the same random intercept effects structure, but different residual variance structures. The null model, which assumed equal residual variance in acceleration between winged and wingless states, was compared to a heterogeneous-variance model that allowed residual variance to differ by wing status using a likelihood-ratio test. Models were fit using lme() function in the ‘nlme’ package [29].

### Species Comparisons

For comparison of tandem running behavior across different species, we observed tandem runs of *M. nervosus* in a standardized method used in previous studies [30,31]. We introduced one female dealate and one male dealate into the arena after 30 minutes of isolation. Both the female and male were marked with one colored dot of paint (PX-20; Mitsubishi) on the abdomen to distinguish sex identities. The observation lasted for 30 minutes after the introduction. In addition to *M. nervosus* dealate pairs, we collected dealates from a single colony of each species and recorded tandem runs of 6 pairs of *Amitermes darwini*, 10 pairs of *Amitermes parvus*, 2 pairs of *Macrognathotermes errator*, 9 pairs of *Macrognathotermes sunteri*, 9 pairs of *Nasutitermes graveolus*, and 2 pairs of *Tumulitermes hastilis* in the same area using the same method as *M. nervosus*. We also collected 20 pairs of *Coptotermes lacteus* from a single colony in Martinsville, NSW, Australia. Information about these colonies is summarized in Table S1. Including *M. nervosus*, we identified species based on the morphological features of soldiers and alates and the distributions, by referring to the Atlas of Living Australia [32] and taxonomic papers [33–35]. All samples are stored at the University of Sydney for future reference.

We analyzed trajectories of these tandem pairs using SLEAP with the same pipeline described above. A model was initialized and trained from scratch to obtain trajectories of *M. nervosus* dealate pairs. We labelled 68 frames in total for *M. nervosus* dealate pairs. This model was then used to pretrain a global model to obtain the trajectories of the seven species used for the comparative analysis. We labeled 210 frames in total to develop our global model. We used the same SLEAP tracking pipeline described above (U-net encoder–decoder architecture with EfficientNet-B0, IOU similarity method, Hungarian matching, and linear interpolation with median smoothing) to infer termite tracks across frames. Pose estimation data from SLEAP were then converted to HDF5 files.

All trajectories were preprocessed (e.g., scaling, down-sampling, body length measurement) and analyzed as described above for comparison of *M. nervosus* pairs before and after wing shed. To enable comparisons across species of different sizes, we standardized movement speed by body length per second. We used a linear model to evaluate differences in running speed during tandem across species using the lm() function in base R. We used Cox proportional hazards models to evaluate differences in tandem and separation duration across species using the coxph() function in the ‘survival’ package in R [36]. After all models were fit, we conducted a post-hoc Dunnett’s test with *M. nervosus* set as the reference level using the glht() function from the ‘multcomp’ package in R [37]. Colony was not included as a random effect for species comparisons because colony variation did not explain additional variation beyond species.

## Results

### Males alter winged female behavior

When paired with a male, 24 out of 36 *M. nervosus* females shed their wings. All wing-shedding events happened after some period of tandem running, with a median time to wing shed of 290 seconds (Fig. 2A). Some females shed their wings in the middle of tandem runs, and other females shed their wings upon separation and resumed the tandem run after re-encounters. In contrast, none of the unpaired females (0/30) shed their wings (Fig. 2A). The movement speed of female termites changed depending on the presence or absence of a male conspecific (Fig. 2B). Movement speed of females in the presence of males was approximately 2.5 times as fast (14.48 mm/s ± 0.98) relative to females in the absence of a male conspecific 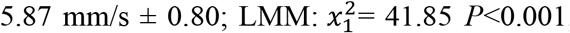, Fig. 2B).

**Figure 2.**
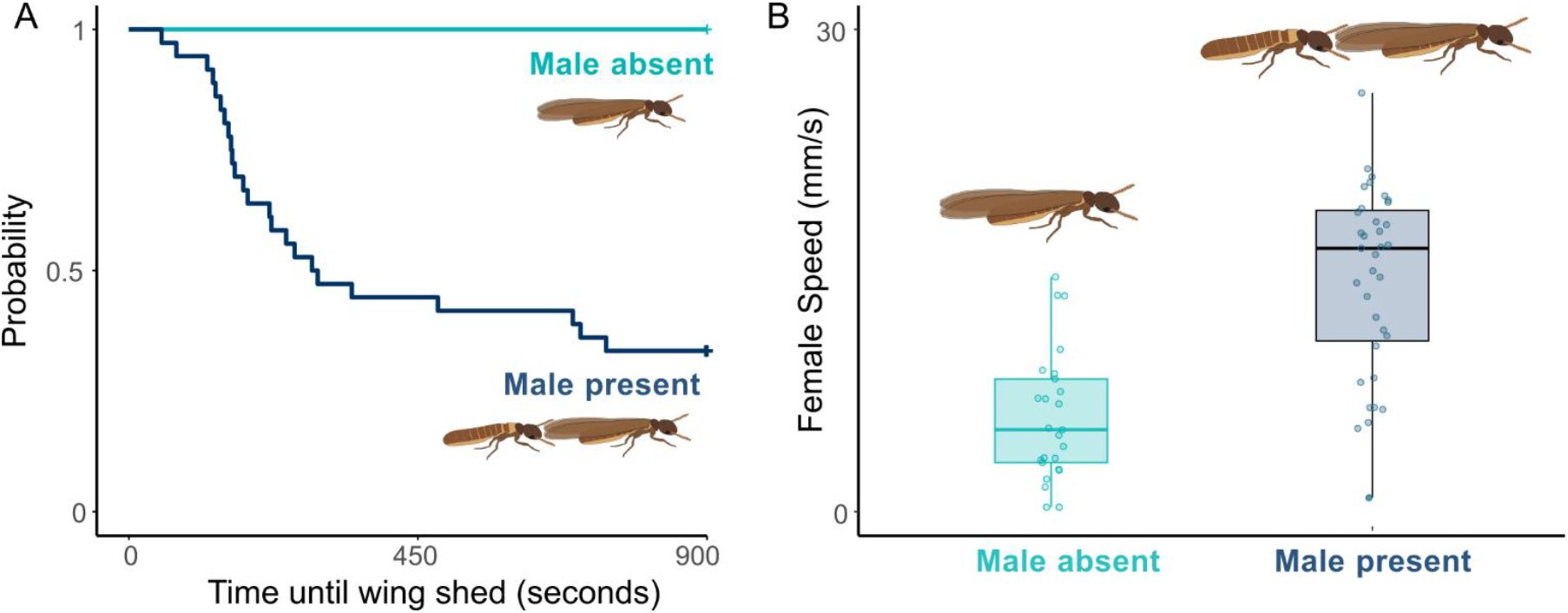
Comparison of A) Probability of female leaders shedding wings. + indicates censored data due to the end of observations. B) female tandem speed in the presence or absence of a male conspecific. In the boxplot, the central line represents the median, box edges represent first and third quartiles, and whiskers extend to 1.5 times the interquartile range for the upper and lower quartiles.

### The presence of wings impairs tandem coordination

The presence of wings did not change overall tandem running speed, although movement speed before wing-shedding displayed larger fluctuations. The model that allowed residual variance to differ by wing status provided a significantly better fit than the homogeneous model to explain the variance of acceleration between winged and wingless states (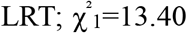, *P* = 0.0003; Fig. S1). Thus, acceleration was more variable prior to wing shedding, with greater frame-to-frame fluctuations in movement speed prior to wing shedding.

We found that tandem runs were much more stable in pairs in which females had shed their wings, compared to those in which females retained their wings (Cox mixed-model: 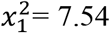, *P* = 0.006; Fig. 3A). Similarly, unwinged pairs required less time to re-establish contact following separation, compared with pairs with winged females, although the difference was not statistically significant, perhaps due to small sample sizes (we observed 16 separation events across 8 winged pairs and 13 separation events across 5 wingless pairs; mixed-effects 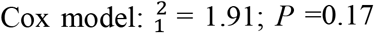, Fig. 3B).

**Figure 3.**
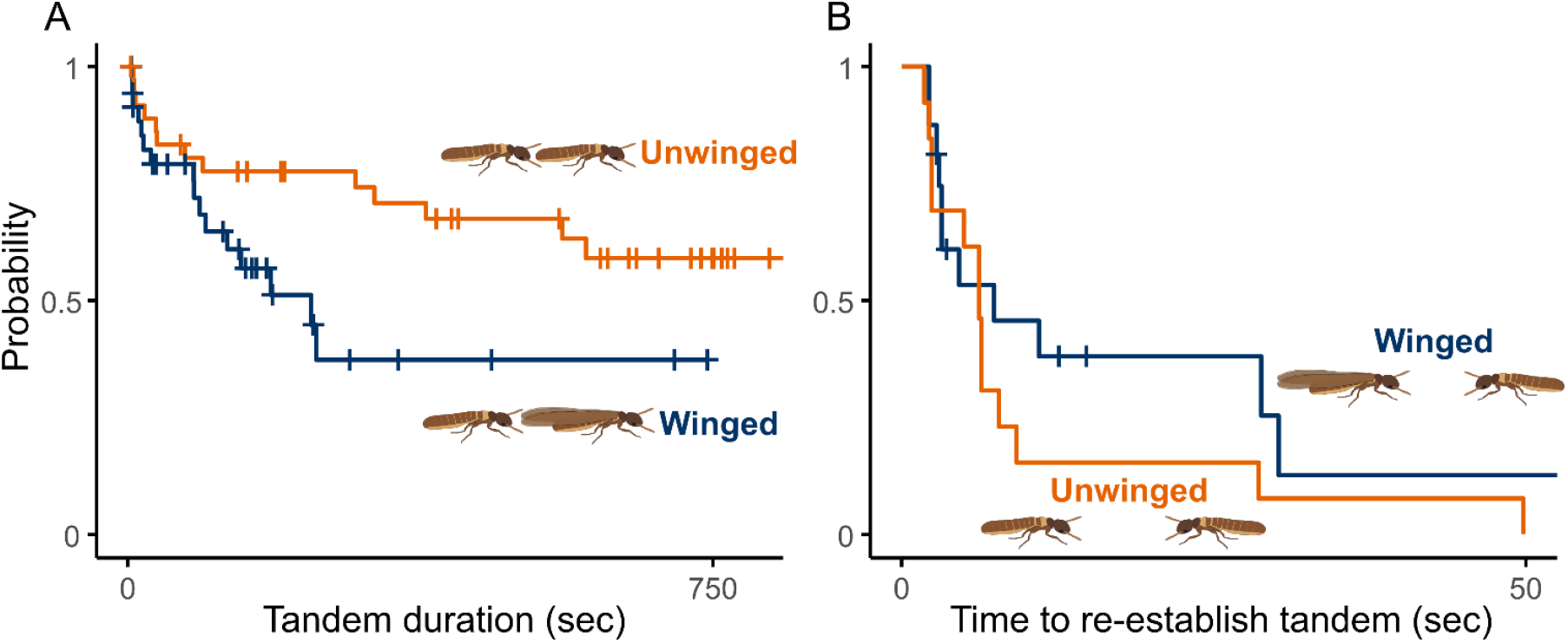
The effect of wings on tandem in *M. nervosus* pairs. **A**) Comparison of the duration of the tandem running event until separation, before and after wingshed. **B)** Comparison of the duration of separation before reuniting between winged and wingless pairs. Note that the x-axis for panel B was cut at 50 seconds, but there was one instance of separation for the winged pairs that extended to 580 seconds. + indicates censored data due to the end of observations.

### 3.3. Comparisons of tandem stability across sympatric termite species

Tandem running speed was different between species (LM, *F*_7,86_ = 18.58, *P* < 0.001, adj. R^2^ = 0.57). Compared to the reference species *M. nervosus*, most species exhibited significantly lower tandem speeds (Fig. 4A; Table S1). Post-hoc Dunnett’s tests confirmed that *Coptotermes lacteus, Tumulitermes hastilis, Nasutitermes graveolus, Macrognathotermes sunteri, Macrognathotermes errator*, and *Amitermes darwini* had significantly reduced tandem speeds relative to *M. nervosus* (Dunnett’s test; *P* < 0.05), with the greatest reduction observed in *Tumulitermes hastilis* (estimate = –2.22 body lengths/s, *P* < 0.001, Table S1). However, the species *Amitermes parvus* did not significantly differ from *M. nervosus* and had a slightly greater tandem speed relative to *M. nervosus* (estimate = 0.30 body lengths/s, *P* = 0.81, Table S1).

**Figure 4.**
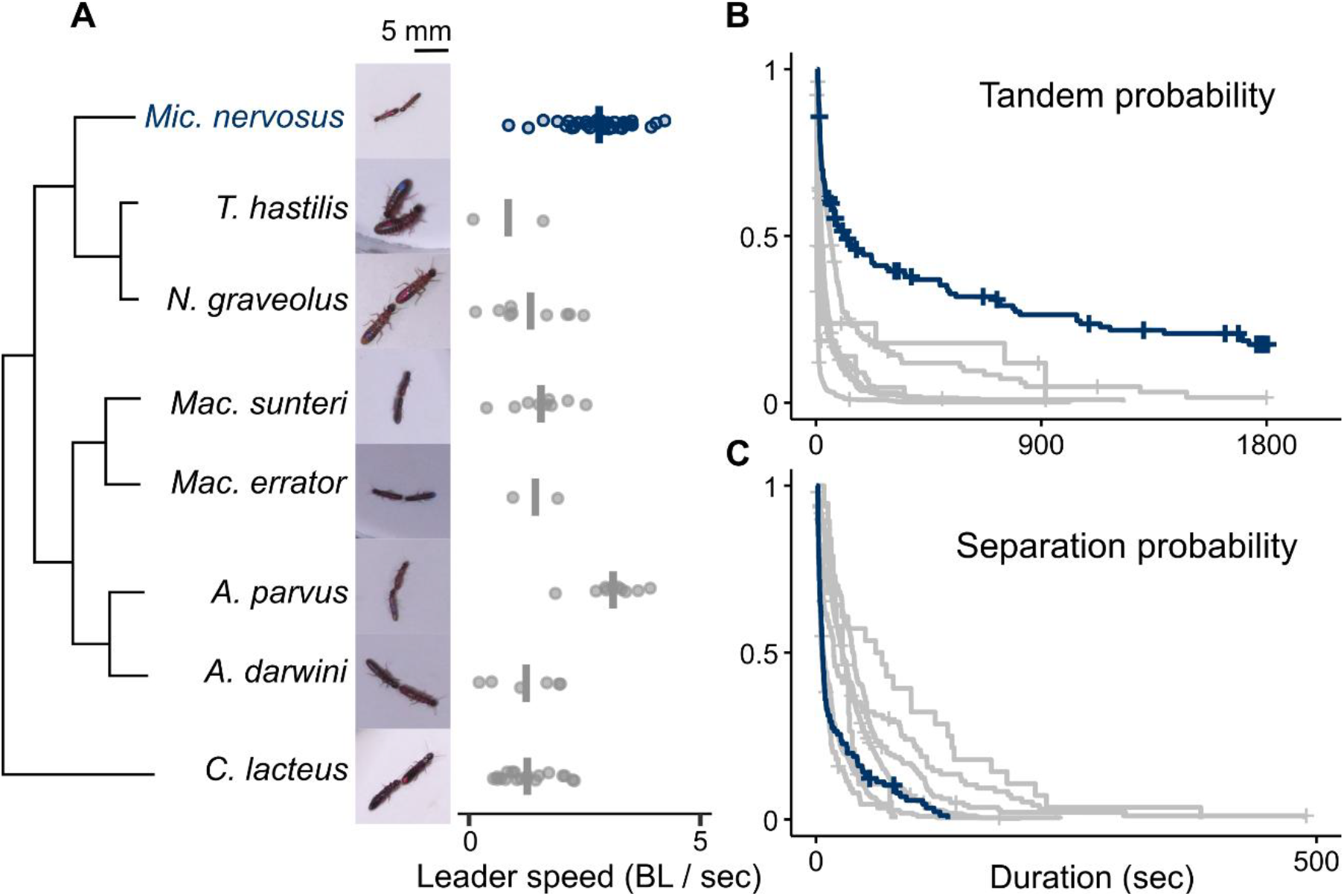
Comparison of tandem running across sympatric species after accounting for differences in body size. A) Tandem speed comparison. The vertical lines denote mean speed. The cladogram is simplified from Hellemans et al. (2024) [38]. B) Comparison of the duration of the tandem running event until separation for each species. C) Comparison of the duration of separation events for each species. *Microcerotermes nervosus* is colored blue for reference.

Tandem running duration also differed between species (Fig. 4B, 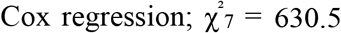, *P* < 0.001). Post hoc Dunnett’s comparisons with *M. nervosus* as the reference group indicated that tandem durations were significantly shorter in all other species (*P* < 0.001, Fig. 4B). Median tandem durations varied substantially among the species sympatric to *M. nervosus*, ranging from just 3.2 seconds in *M. errator* to 57.6 seconds in *A. parvus*. In contrast, the median tandem duration of our focal species, *M. nervosus*, was markedly longer (106.8 seconds), greatly exceeding that of all other species compared. Note that neither tandem speed nor tandem stability was significantly related to female body size across species (Fig. S3).

Separation duration also varied significantly between species (Fig. 4C; 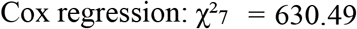, *P* < 0.001). Using *M. nervosus* as the reference, post hoc Dunnett’s tests showed that separation durations were significantly longer in most tested species (*P* < 0.001). Specifically, it took longer for *A. darwini, C. lacteus, M. errator*, and *M. sunteri* to reunite with their partner than *M. nervosus*. Separation durations of *A. parvus* were significantly shorter than those of *M. nervosus*.

## Discussion

Our findings demonstrate that mate pairing behaviors of *M. nervosus* deviate from the typical neoisopteran precopulatory sequence, as a result of engagement in tandem running prior to wing shedding. In most termites, dealation has been considered a prerequisite behavioral event preceding tandem running because wings can hinder mobility and communication [13,14]. In *M. nervosus*, however, tandem running itself could be the prerequisite for female dealation (Fig. 2). This reordering represents a novel mechanism of behavioral diversification. Previous work has emphasized that diversification in mating sequences can occur through the addition, deletion, or modification of components, including changes in their duration or intensity [39–44]. Our results, on the other hand, show that diversification can also arise through the reshuffling of preexisting elements, expanding our understanding of the pathways for the evolution of behavioral sequences. Our results also emphasize why dealation is typically conserved as a prerequisite to tandem running in other termites. During tandem running, the following males rely on antennal and palpal contact with the leading female abdomen to maintain physical contact [13,45]. Wings of females interfere with this tactile signaling essential for pair coordination. In agreement with this, we found that tandem coordination was less stable in winged vs unwinged *M. nervosus* (Fig. 3A). Similarly, wings might prolong the time required for pairs to reunite after separation, although this difference was not statistically supported (Fig. 3B). Although such “signal jamming” is widely observed in other systems, interference is usually due to external factors, e.g., acoustic masking in birds and marine mammals, where background noise reduces the effective range of signals [46,47], or speech impediments in humans wearing retainers, which impair certain articulations [48,49]. In termites, such interference originates from the behavioral sequence of mating behavior, where pairs need to perform walking movement coordination after the dispersal flight. This context likely explains why males of most species do not attempt to follow females before dealation.

Comparisons of sympatric species provide additional explanation for why *M. nervosus* has evolved this flexibility. During tandem running, *M. nervosus* exhibited much more stable movement coordination than other species (Fig. 4). Even with winged females, *M. nervosus* exhibited tandem stability greater than or comparable to that of all tested sympatric species without wings (Fig. S2). This inherent high ability to coordinate movement may have allowed this species to perform tandem running with the interference of wings. Which ecological pressure facilitated this behavioral reordering? One possible candidate is termite nesting ecology, where nest site selection by delates determines the location of subsequent colony growth. Compared with other species, *M. nervosus* is clearly less selective when it comes to choosing nest sites. This species constructs small mound nests across northern Australia, often being the most abundant mound builders in the area (see Fig. 1A for the high density of mound nests in the studied area) [50]. Such generalist nesting habits may reduce the selective pressure on immediate nest discovery; instead, *M. nervosus* may prioritize securing a partner even before wing-shedding. This finding highlights the importance of a holistic understanding of life history traits beyond mating to understand the evolution of sequential behaviors.

Proximately, what internal processes can enable the reversed order of mating sequence in this species? Our data show that females do not shed their wings in isolation but do so within minutes of initiating a tandem pair with a male conspecific. This minutes-level latency period between initiation of tandem and wing shedding matches the timescale of rapid neuroendocrine responses to stimuli found in other invertebrates [51–54]. One possibility is that tactile stimulation from the male’s antennae and palps triggers hormonal cascades of biogenic amines that promote dealation. Alternatively, repeated mechanosensory input during tandem may drive neural activity toward a threshold that releases the wing-shedding motor action (e.g., ramp-to-threshold mechanism), akin to neuronal control of mating sequence order in *Drosophila melanogaster* [4,5]. Future work combining behavioral assays with neurophysiological or pharmacological manipulations will disentangle these mechanistic hypotheses.

In summary, we identified a novel route to behavioral diversification through the reordering of events in a strongly integrated behavioral sequence. By showing that tandem running can precede and facilitate dealation, *M. nervosus* illustrates that modular components capable of reorganization can exist even in functionally conserved behavioral architectures. More broadly, it raises questions as to how widespread shifts in behavioral order occur across taxa, what selective mechanisms mediate them, and whether such changes can contribute to reproductive isolation and speciation.

## Contributions

EC: Software, Data curation, Validation, Formal analysis, Visualization, Writing – Original Draft. TK: Investigation, Writing – Review & Editing.

NL: Resources, Writing – Review & Editing.

NM: Conceptualization, Investigation, Methodology, Validation, Supervision, Funding acquisition, Writing – Review & Editing.

## Acknowledgements

We thank Kensei Kikuchi for helping during the sampling of termites, and Aoi Mizumoto for helping during the video processing. The University of Sydney kindly provided accommodation and laboratory space at the Crommelin Research Station.

