## Supplementary material for "Diversification of pre-mating behaviors through temporal reordering of components": Table S1, Figure S1-3

This file includes

Table S1

Figure S1-3

**Table S1.** Collection localities, dates, and colony codes of seven sympatric termite species used in comparative analyses with *Microcerotermes nervosus*.

| Genus | Species | Date | Latitude | Longtitude |
|---|---|---|---|---|
| *Amitermes* | *darwini* | 10/31/2023 | -12.8020 | 131.1016 |
| *Amitermes* | *parvus* | 10/31/2023 | -12.8020 | 131.1016 |
| *Coptotermes* | *lacteus* | 10/22/2023 | -33.0672 | 151.3326 |
| *Macrognathotermes* | *errator* | 10/28/2023 | -12.4785 | 131.0302 |
| *Macrognathotermes* | *sunteri* | 10/29/2023 | -12.4607 | 131.0345 |
| *Nasutitermes* | *gravelous* | 11/1/2023 | -12.4384 | 130.8485 |
| *Tumulitermes* | *hastilis* | 10/28/2024 | -12.4769 | 131.0295 |

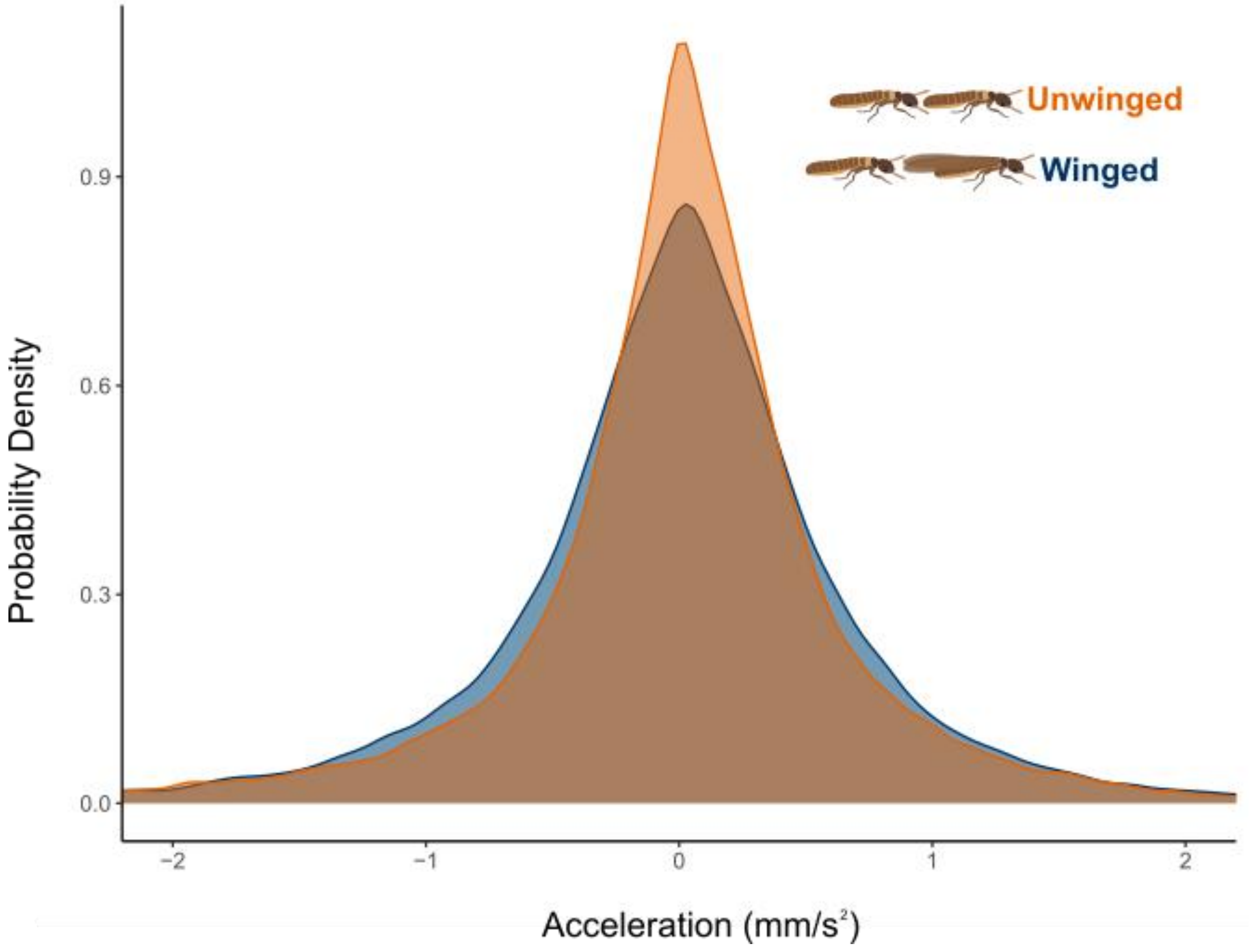


**Figure S1.** Probability density distributions of acceleration during tandem running for *M. nervosus* alates before wing shedding and after wing shedding. Both states show a peak at low acceleration values near zero, but unwinged individuals exhibit a narrower distribution with reduced variance compared to winged individuals.

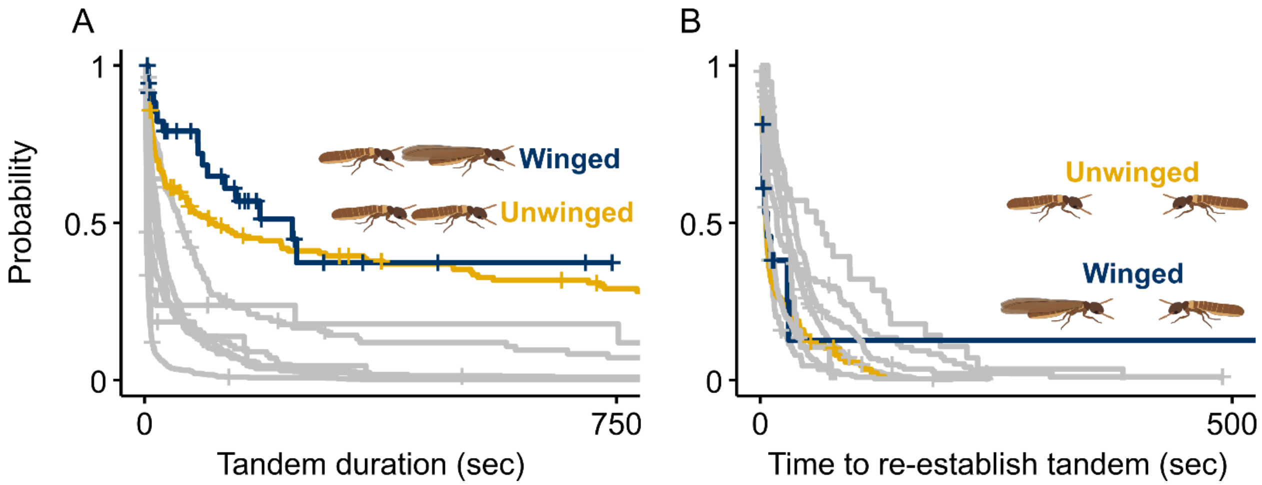


**Figure S2.** Comparison of A) the duration of the tandem running event until separation for each species and B) duration of separation events for each species. Winged pairs of *M. nervosus* were included here to provide a comparison of their tandem stability prior to wing shedding (Fig. 3).

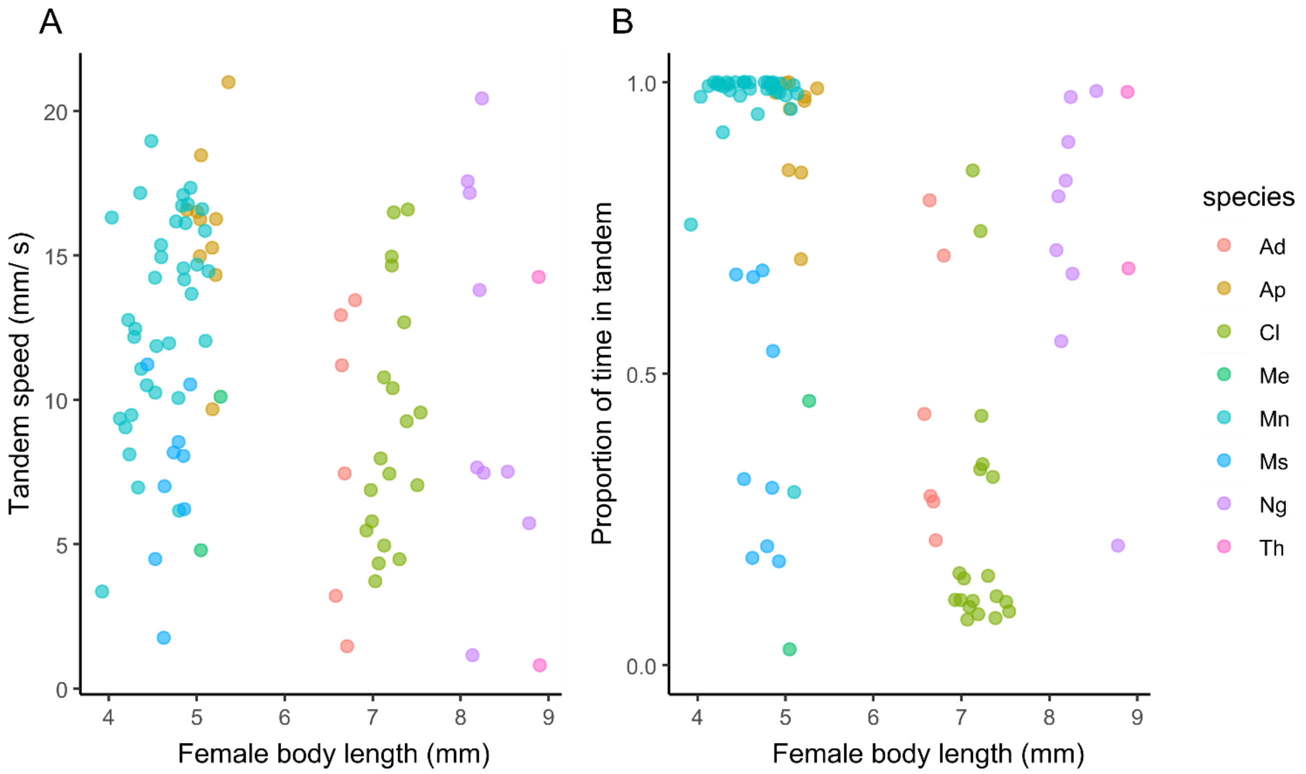


**Figure S3.** The relationship between female body size and A) tandem speed and B) tandem duration within the 30-minute period. For analyses of tandem duration, proportion data were logit-transformed. We fit linear mixed-effects models using the lmer() function in the "lme4" package, with female body length as a continuous covariate, random intercepts for species, and random slopes that account for how speed changes with body size within species. This structure allows each species to have its own mean tandem speed or duration and its own relationship between female body length and tandem behavior, thereby capturing among-species differences while accounting for within-species variation in body size. We found that neither tandem speed nor the proportion of time spent in tandem was significantly related to female body size across species. (LMMs; speed: $\chi^2_1 = 0.0002$, $P = 0.99$; tandem duration: $\chi^2_1 = 0.07$, $P = 0.79$).
